# Guanine content in the CTCF motif encodes a vulnerability to persistent, alcohol-induced redistribution

**DOI:** 10.64898/2026.10.14.759128

**Authors:** Ben Nolan, Joynob A. Puspo, Iesha M. Querry-Peterson, Stephanie Maurina, Rebecca A. Wright, Allyson Schmitz, Lauren E. Jacobsen, Coralmarie Ortiz, Timothy E. Reznicek, Alexander Ethell, Hannah L. Harris, Christopher T. Cummings, M. Jordan Rowley

## Abstract

Alcohol consumption is a leading contributor to the global disease burden, yet its molecular consequences remain incompletely understood. Ethanol’s first metabolite, acetaldehyde, modifies guanine to form DNA adducts and DNA-protein crosslinks that block elongating RNA polymerase II. Because acetaldehyde acts specifically on guanine, we hypothesized that chromatin features would be disrupted in a sequence-dependent rather than stochastic manner. Here we confirm that alcohol exposure displaces elongating RNA polymerase II (RNAPIIs2p) from gene bodies, relocating to promoters. In addition to this confirmation, we found that RNAPIIs2p also relocates to discrete guanine-rich intergenic CTCF sites. Loci with high guanine content at the CTCF motif’s zinc finger 5 (ZF5) binding position showed the largest increase in both CTCF and cohesin (RAD21), with CTCF occupancy changes persisting even six days after alcohol withdrawal. Surprisingly, using Hi-C, we found that these changes to CTCF and cohesin do not impact the pronounced structural CTCF loops, but are instead primarily associated with altered guanine-rich enhancer-promoter contacts that connect to lasting changes to gene expression. Beyond our cell-line model, the same sequence rule accounts for altered CTCF occupancy in a mouse starvation (metabolic stress) model and in the placentae after paternal alcohol exposure. Together these findings link alcohol exposure to sequence-directed changes to chromatin whose rules are conserved in alcohol-associated disease.

**GRAPHICAL ABSTRACT:** 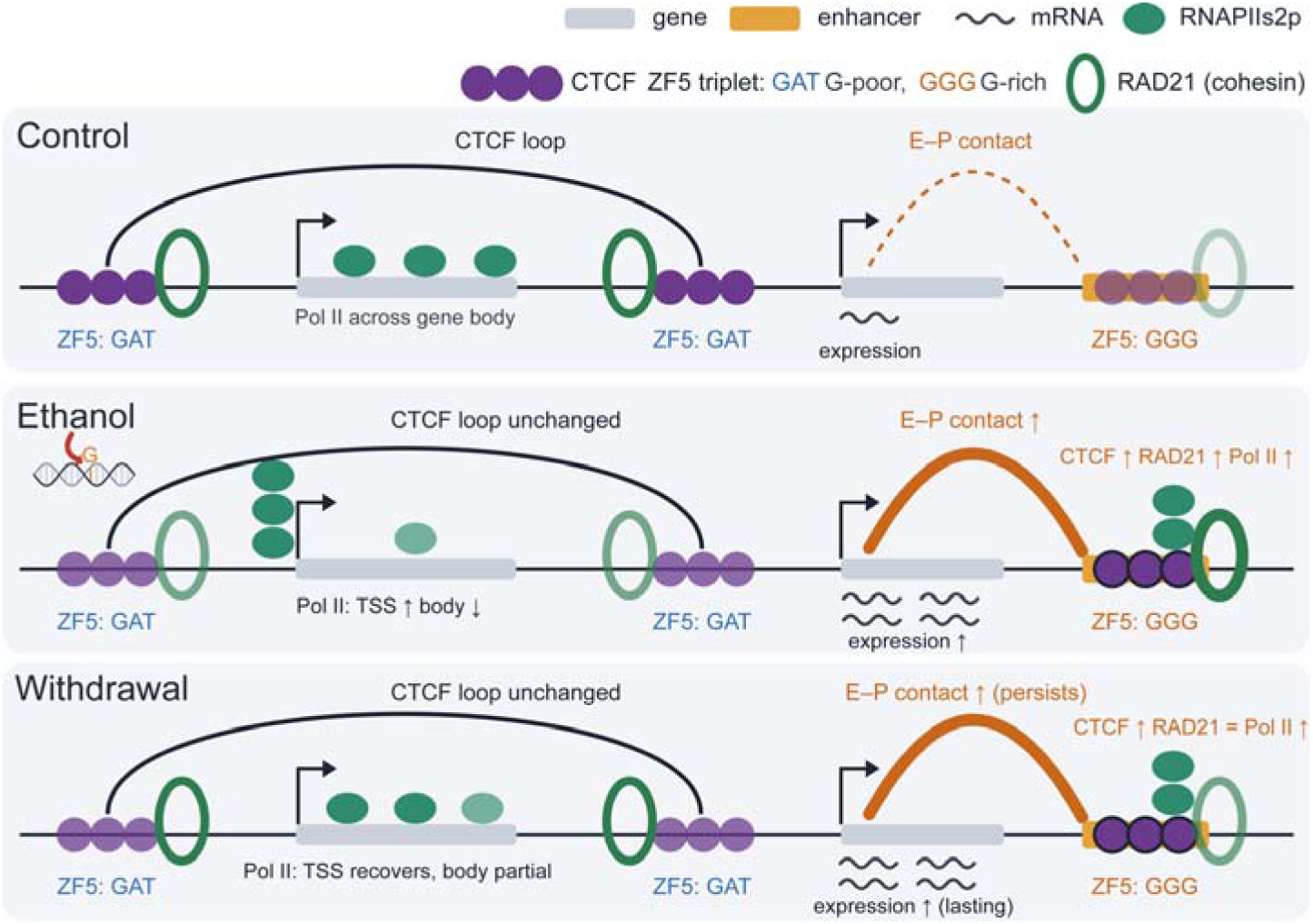

## INTRODUCTION

Alcohol is a widely consumed toxic substance that was responsible for 4.7% of all deaths globally in 2019^1^. Its first cellular metabolite, acetaldehyde, is a highly reactive electrophilic compound that establishes covalent bonds between guanines as inter- and intrastrand crosslinks, and between guanine and proximal proteins as DNA-protein crosslinks (DPC)^2–4^. These crosslink-retained proteins represent a large physical block to processes that track along DNA^3,4^. Clinically, deficiencies in crosslink repair cause severe disease, including Fanconi anemia, Ruijs-Aalfs syndrome, and AMeD syndrome^5–9^. Yet, while acetaldehyde is a known guanine crosslinker, the broader consequences for chromatin structure have not been elucidated.

While little is known about the relationship between alcohol exposure and chromatin, more is known about how cells handle damage from other reactive events, such as the crosslinks and adducts induced by UV radiation, formaldehyde and oxidative stress^10–12^. These Transcription-Blocking Lesions (TBL) stall RNA polymerase II (RNAPII), which in turn recruits the machinery that removes the lesion and allows transcription to resume^13^. This transcription-coupled surveillance operates even under normal conditions, because reactive aldehydes are generated continuously by endogenous metabolism^14^. In these models, damage is generally treated as stochastic, with lesion burden scaling with the size of the transcribed region rather than with any particular sequence^10,12,15^. Acetaldehyde from alcohol metabolism, therefore, does not act on a naïve genome; instead, it adds to a standing burden of aldehyde-driven crosslinks that cells are already shaped by and equipped to resolve, up to a point. What alcohol changes is the scale of that burden, and whether the repair machinery can keep pace.

The architectural protein CTCF organizes the genome into thousands of loops and loop domains^16,17^. In combination with the cohesin complex, chromatin is extruded into structural loops which appear as intense punctate signal in Hi-C maps of chromatin architecture^18–20^. Apart from its role with cohesin in forming these canonical ‘CTCF loops’, CTCF is also present at thousands of additional sites throughout the genome which are thought to facilitate enhancer-promoter contacts^21^. CTCF recognizes its DNA binding loci through an array of zinc fingers, some of which contacting a nucleotide triplet that together comprise an extended, guanine-rich core motif^22^. The guanine content at motif core has been shown to correlate positively with CTCF binding strength^23^. We hypothesized that the high guanine content characteristic of the CTCF motif makes it susceptible to acetaldehyde^24–26^. However, it is yet to be established whether alcohol preferentially perturbs DNA binding proteins such as CTCF which is reliant on guanine-rich motifs, and whether the natural sequence variation between CTCF binding sites predicts enhanced susceptibility at specific loci, and whether downstream genome organization and gene expression are disrupted as a result.

Here we use an exposure-and-withdrawal model of binge-level ethanol exposure in human cells to determine how ethanol reshapes the chromatin landscape and whether the changes persist after exposure ends. Profiling RNAPII, CTCF, and RAD21 occupancy, alongside 3D contacts by Hi-C, we show that alcohol redistributes the occupancy of elongating RNAPII, CTCF, and cohesin. Enhancer-promoter interactions, but not the pronounced structural CTCF-CTCF loops, are disrupted with corresponding alterations to gene expression. The majority of these changes remain even six days after alcohol withdrawal, indicative of a persistent change at the chromatin level. We found that the impact of alcohol on CTCF is encoded in the motif itself, particularly corresponding to the guanine content within the binding site contacted by zinc finger 5. Furthermore, we observe that these rules explain previously demonstrated CTCF changes in a mouse starvation (metabolic stress) model and in the placentae of offspring from alcohol-exposed sires. Together these results link alcohol exposure to persistent, sequence-encoded changes to chromatin architecture, with relevance to the diseases associated with its consumption.

## RESULTS

### Alcohol induces gene expression changes that persist 6 days after withdrawal

Cells were exposed to fresh media containing 200 mg/dL ethanol daily for 6 days, a concentration used previously to model binge-level exposure^27,28^ (Fig. 1A). Control cells received alcohol-free media on the same schedule, and the withdrawal condition included the 6-day ethanol treatment along with a further 6 days of alcohol-free media to test recovery (Fig. 1A). Neither cell count nor cell cycle distribution differed between control and ethanol-treated cells over the treatment period (Fig. 1B, Table S1), consistent with the sub-cytotoxic dose used in alcohol-exposure models ^29,30^.

**Figure 1:**
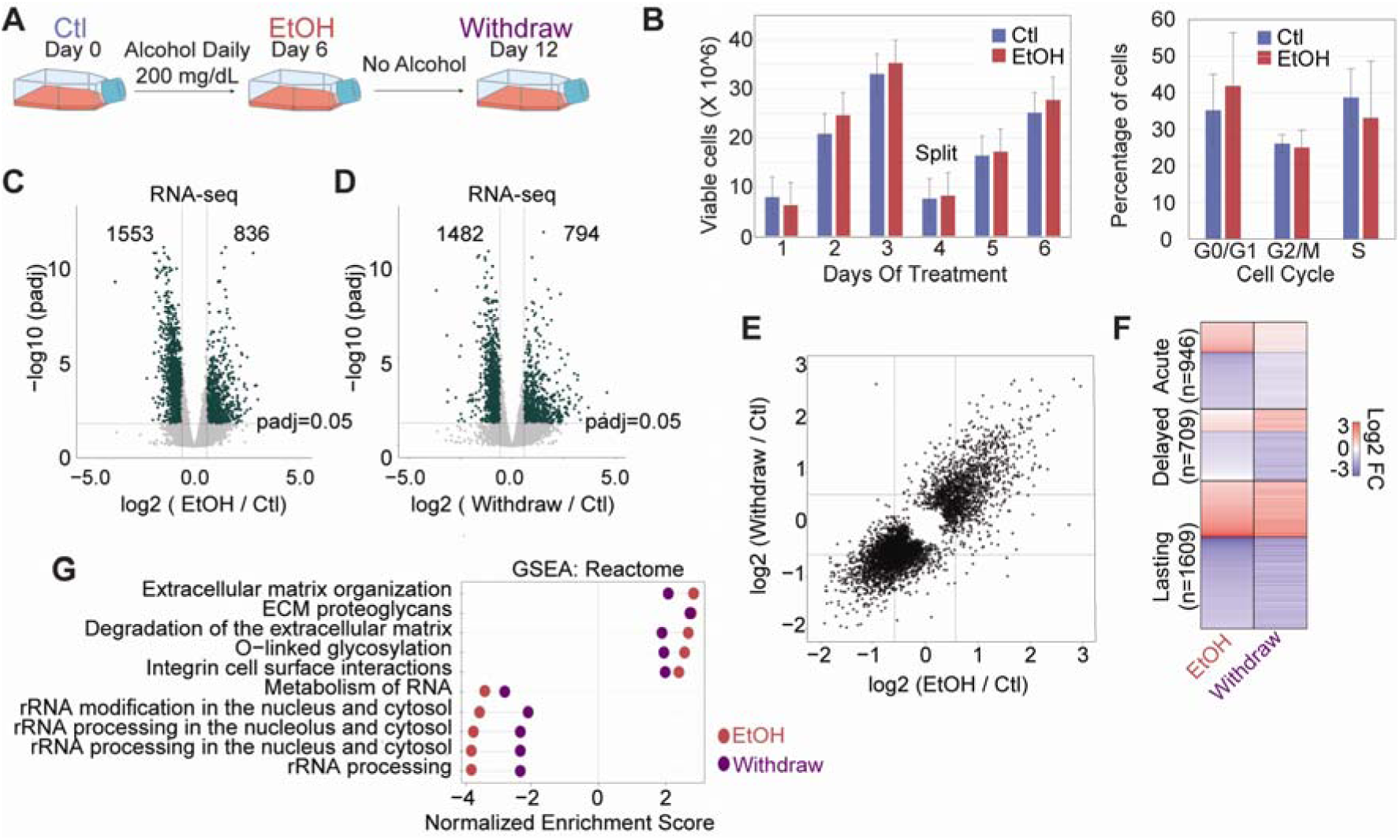
A) Experimental design where cells were maintained in alcohol-free media (Ctl), exposed to 200 mg/dL ethanol for 6 days (EtOH), or exposed for 6 days followed by 6 days of an alcohol-free period (Withdraw). These three conditions apply to every assay in this study unless otherwise stated. B) Daily cell count (left) and cell cycle count (right) in Ctl and EtOH conditions. C&D) Gene expression distribution after ethanol (C) or withdrawal (D) with dotted line and counts representing a padj< 0.05, 1.5-fold change cutoff. E) Scatter plot of the expression log2 fold changes (vs Control) for Ethanol (x-axis) compared to Withdrawal (y-axis) across all differentially expressed genes (p-adjusted < 0.05). Dashed lines indicate a +/- 1.5 fold change in expression. F) Heatmap of differentially expressed genes (1.5-fold change, p-adj < 0.05) categorized by changes in both EtOH and Withdraw (Lasting), unchanged in EtOH with changes in Withdraw (Delayed), or changes in EtOH followed by a recovery in Withdraw (Acute). G) Gene Set Enrichment Analysis (GSEA: Reactome) comparing pathways associated with gene expression changes after EtOH and Withdraw.

To define ethanol’s impact on gene expression in our model, we profiled genome-wide gene expression by RNA-seq (Fig. S1). Ethanol exposure induced widespread changes in steady-state gene expression, with 2,555 genes differentially expressed relative to control (1,553 down, 836 up; p-adj < 0.05, |fold change| > 1.5; Fig. 1C). The majority of differentially expressed genes (1,609) showed persistent dysregulation, mainly downregulation, even 6 days after withdrawal (Fig. 1D-F). Other notable categories included 946 genes that partially recovered (acute response) and 791 genes that only became differentially expressed after alcohol withdrawal (delayed response) (Fig. 1F). Independent of these groupings, gene set enrichment analysis also demonstrated that the top 10 pathways altered by ethanol remain altered after 6-day withdrawal from ethanol (Fig. 1G), with ribosomal RNA processing and biogenesis downregulated and extracellular matrix organization upregulated. The suppression of ribosome biogenesis is notable, as it has been reported in cellular and animal models of alcohol teratogenicity and proposed as a mechanism in fetal alcohol spectrum disorder ^31–35^. Thus, despite unchanged proliferation and cell cycle progression, sub-cytotoxic ethanol exposure (binge-level) produces substantial gene expression changes that persist days after the exposure ends.

### Alcohol stalls RNA polymerase II at promoters and CTCF sites at the expense of gene-body elongation

The persistence of expression changes, even across multiple cell divisions in the withdrawal condition, led us to interrogate a possible molecular memory of ethanol exposure. Therefore, we asked if there are underlying changes to chromatin. First, we mapped the elongating form of RNAPII (serine 2 phosphorylation; RNAPIIs2p) by CUT&Tag. During ethanol exposure, RNAPIIs2p was increased at promoters while decreasing over the gene body, across all expression groups (Fig. 2A). During withdrawal, RNAPII occupancy at the TSS recovers, but only partial recovery was observed at the gene body (Fig. 2A).

**Figure 2:**
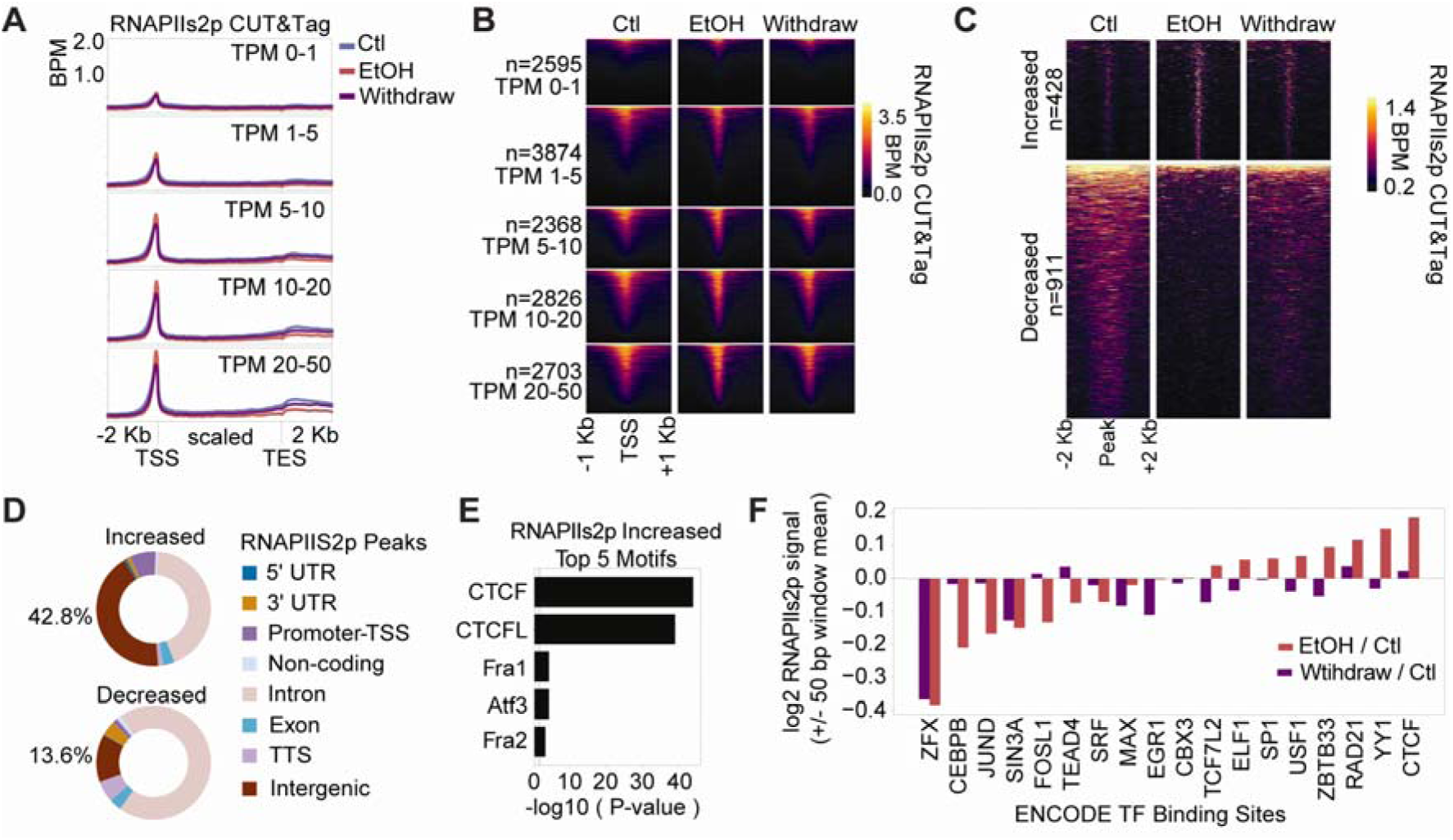
A) Average RNA polymerase II serine 2 phosphorylation (RNAPIIs2p) CUT&Tag signal across scaled genes in control, ethanol treated, and withdrawal cells. Genes are grouped by control TPM expression values as indicated. B) Heatmap of RNAPIIs2p CUT&Tag signal (BPM) at the TSS in the same TPM groups. C) RNAPIIs2p signal (BPM) at differential peaks after EtOH exposure. D) Genomic regions associated with RNAPIIs2p differential peaks. E) Top 5 known motifs at differentially increased RNAPIIs2p peaks. F) RNAPIIs2p mean BPM signal enrichment in both EtOH and withdraw conditions across 18 transcription factor binding sites.

Heatmaps of the CUT&Tag signal emphasize this pattern of concentrated RNAPIIs2p signal directly at the TSS coinciding with decreases in the proximal diffuse gene body signal after ethanol exposure (Fig. 2B). Differential peak analysis identified 1,339 differential RNAPIIs2p peaks following ethanol exposure, revealing that decreased peaks were characteristically broad and ∼79% intragenic (intronic, exonic, TTS) (Fig. 2C, 2D). In contrast, regions with increased RNAPIIs2p signal were sharp peaks, enriched in intergenic and promoter-TSS regions (Fig. 2C, 2D).

To determine whether increased RNAPIIs2p at intergenic sites corresponds to specific transcription factor binding sites, we performed motif analysis which revealed enrichment of the CTCF motif (Fig. 2E). In contrast, lost RNAPIIs2p showed a discernibly lower significance of motifs (Fig. S2A). We then measured RNAPIIs2p at the binding sites of 18 transcription factors annotated by ENCODE ^36^ which found that ZFX had the largest decrease in RNAPIIs2p, consistent with its preferential location downstream from the TSS ^37^ (Fig. 2F). Similar to the motif analysis, CTCF occupied sites had the highest increase in RNAPIIs2p signal after ethanol exposure, followed by YY1 and RAD21 which often coincide with CTCF (Fig. 2F, Fig. S2B, Table S2) ^38^. Together, these data reveal an alcohol-induced redistribution of RNA polymerase away from gene bodies and towards CTCF binding sites.

### Alcohol exposure shifts CTCF and RAD21 chromatin occupancy toward guanine-rich sites

As RNAPIIs2p-gained sites matched the CTCF motif (Fig. 2E), we examined how alcohol affects CTCF’s pattern of chromatin occupancy. Given that acetaldehyde targets guanine, and that the CTCF motif is guanine-rich, we further asked whether the guanine content of individual CTCF binding sites determines the response to ethanol. We performed ChIP-seq for CTCF and found that in control cells, occupancy was only weakly correlated with GC content (Spearman = 0.192) (Fig. 3A-B). During ethanol exposure this correlation strengthened (Spearman = 0.292), as CTCF was diminished at low-GC sites and gained at high-GC sites (Fig. 3A-B). As an example, CTCF occupancy is increased after ethanol exposure at the promoter of *RRP1*, a ribosomal RNA processing gene that is itself downregulated after ethanol (Fig. 1E, 3C). Notably, we checked if mutations were induced by 200 mg/dL ethanol exposure but found they occurred at subclonal allele frequencies (Fig. S3A), indicative of no significant mutagenic difference caused by ethanol treatment. For confirmation, we also examined potential SNV candidates and found no relationship with alcohol-sensitive CTCF sites (Fig. S3B) indicating that changes to CTCF occupancy are not attributable to a change in the underlying sequence.

**Figure 3:**
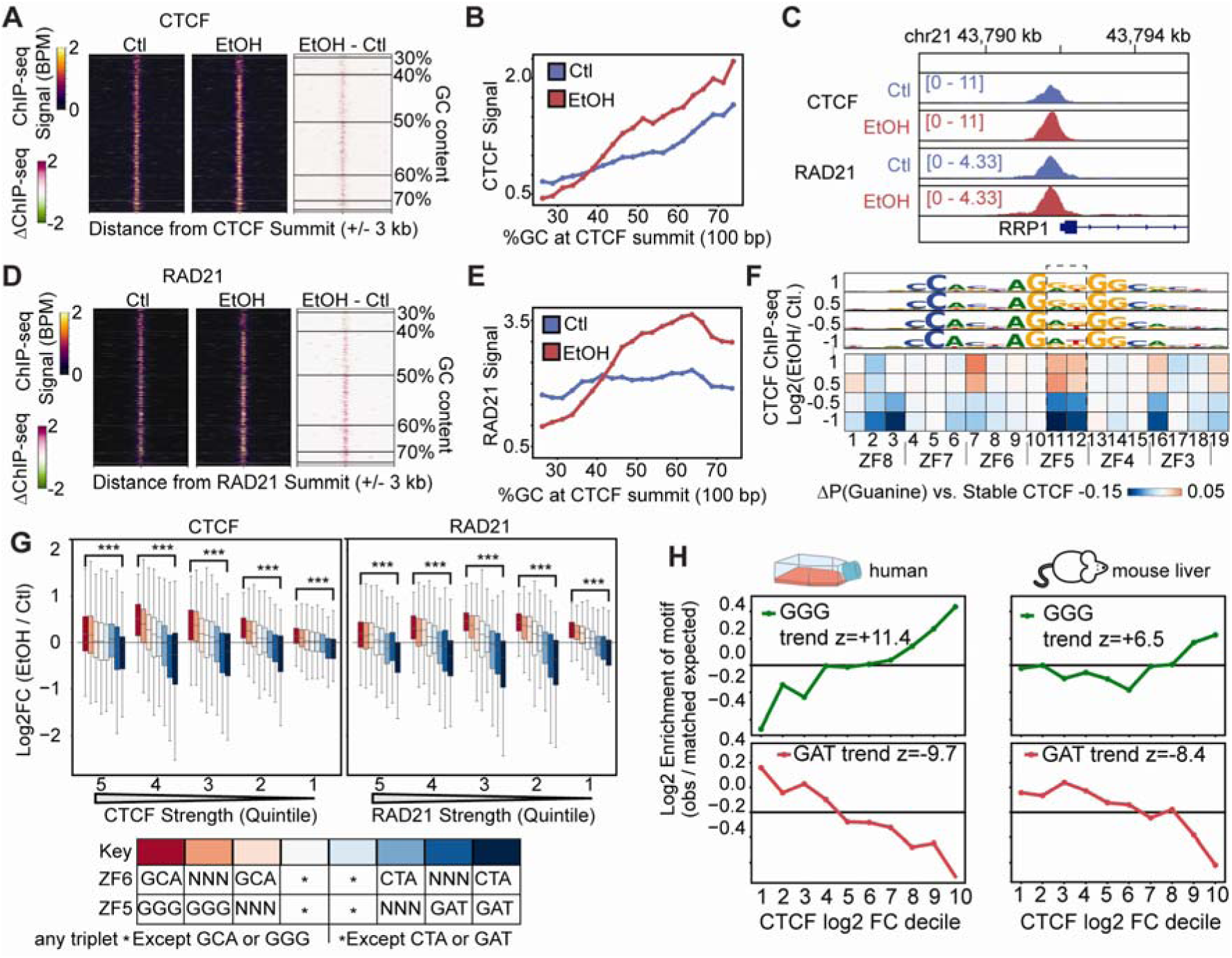
A&D) CTCF (A) and RAD21 (D) BPM signal heatmap in control (Ctl) and ethanol (EtOH) conditions with a differential (EtOH – Ctl) heatmap (right). Sites are ordered by GC content. B&E) Mean CTCF (B) and RAD21 (E) signal ordered by GC content. C) Example locus (RRP1) with an increase in CTCF and RAD21 chromatin occupancy after ethanol exposure. F) CTCF peaks resolved to a single 19 bp motif sequence, categorized by log2 fold change. Heatmap demonstrates guanine probability compared to unchanged peaks, with motif logo for the same groups plotted above. G) CTCF and RAD21 binding strength difference in EtOH condition at CTCF sites with zinc finger 6 and 5 as ‘GCA’, ‘CTA’ or ‘GGG’, ‘GAT’ respectively across binding strength quintiles. *Any triplet excluding the triplets being measured. H) Enrichment of ZF5 ‘GGG’ and ‘GAT’ by CTCF log2 fold change decile in two models: human cells CTCF ChIP-seq after alcohol exposure for 6 days, and publicly available CTCF ChIP-seq data from ad-lib fed and 24-hour starved mouse livers^39^. *** Wilcoxon-rank test < 0.001

ChIP-seq for RAD21, a subunit of cohesin, revealed a similar pattern, with the largest gains at GC-rich loci (Fig. 3D-E). RAD21 levels at CTCF sites did not correlate with GC content in control cells (Spearman = 0.014) but had a dramatic increase in GC correlation during ethanol exposure (Spearman = 0.199) (Fig. 3E).

Considering CTCF redistribution tracked with GC content genome-wide, we reasoned that specific zinc finger binding positions within the motif may drive this effect. Ethanol-induced changes to CTCF occupancy showed a strong preference for sites which had guanine present at motif base-pair positions 7, 11, 12, and 16 (Fig. 3F), corresponding to sites recognized by zinc fingers (ZF) 6, 5, and 3 respectively^22^. Indeed, CTCF sites with ‘GGG’ at ZF5 have a mean increase in occupancy after ethanol exposure for both CTCF and RAD21 independent of baseline binding strength (Fig. 3G).

Because we found bigger increases were associated with guanine at multiple zinc fingers, we asked if their contributions are potentially additive with each other. For example, we found the strongest increases in CTCF occupancy at loci with both GCA and GGG at ZF6 and ZF5 positions respectively (Fig. 3G, dark red), and the strongest decreases at CTCF loci with both CTA and GAT at these positions (Fig. 3G, dark blue). Indeed, across all pairwise combinations of zinc finger triplets, the predicted additive effects and observed combined effects were strongly correlated (*R* = 0.87; Fig. S3C), indicating that CTCF vulnerability to ethanol corresponds to local DNA sequence, with individual zinc fingers acting additively.

To establish whether guanine content at ZF5 predicts CTCF’s response to ethanol, we performed permutation testing for ZF5 ‘GGG’ or ‘GAT’ across deciles of the log2 fold change after ethanol and found a significant enrichment of these motifs in gained and lost sites respectively (Fig. 3H). Furthermore, we asked whether this rule is specific to ethanol or reflects a more general property of CTCF under metabolic stress by reanalyzing published CTCF ChIP-seq data from mice in starved vs. fed conditions, a model of metabolic stress ^39^. Notably, in this previous study, they found that CTCF occupancy on chromatin was overall higher in fed conditions (representing higher levels of metabolically produced aldehydes), which would be consistent with our ethanol response. Applying the same motif-based analysis, we found that ZF5’s GGG was overrepresented, while GAT was underrepresented, at the top gained CTCF sites (Fig. 3H). These results suggest that guanine content at ZF5 governs not only CTCF’s sensitivity to alcohol but also its response to diet-associated metabolically produced aldehydes broadly.

### Persistent CTCF redistribution encodes a memory of alcohol exposure

Gene expression changes persisted six days after ethanol withdrawal (Fig. 1). As such, we asked whether CTCF occupancy showed similar persistence. RNAPIIs2p partially recovered after withdrawal, with incomplete return to baseline occupancy at CTCF sites among other TF binding sites (Fig. 2F). CTCF occupancy, by contrast, remained increased at high-GC sites (Fig. 4A-B), indicating that the alcohol-induced redistribution of CTCF along the GC axis is largely stable on this timescale. Furthermore, the motif features that predicted the ethanol response remained predictive after withdrawal, with zinc-finger triplets still contributing additively (Fig. S4A) and both motif composition and per-position guanine probability continuing to separate CTCF sites by susceptibility (Fig. S4B). Thus, cellular exposure to ethanol induces a persistent change in the CTCF binding landscape that outlasts the exposure (see example in Fig. 4C, CTCF peak 2).

**Figure 4:**
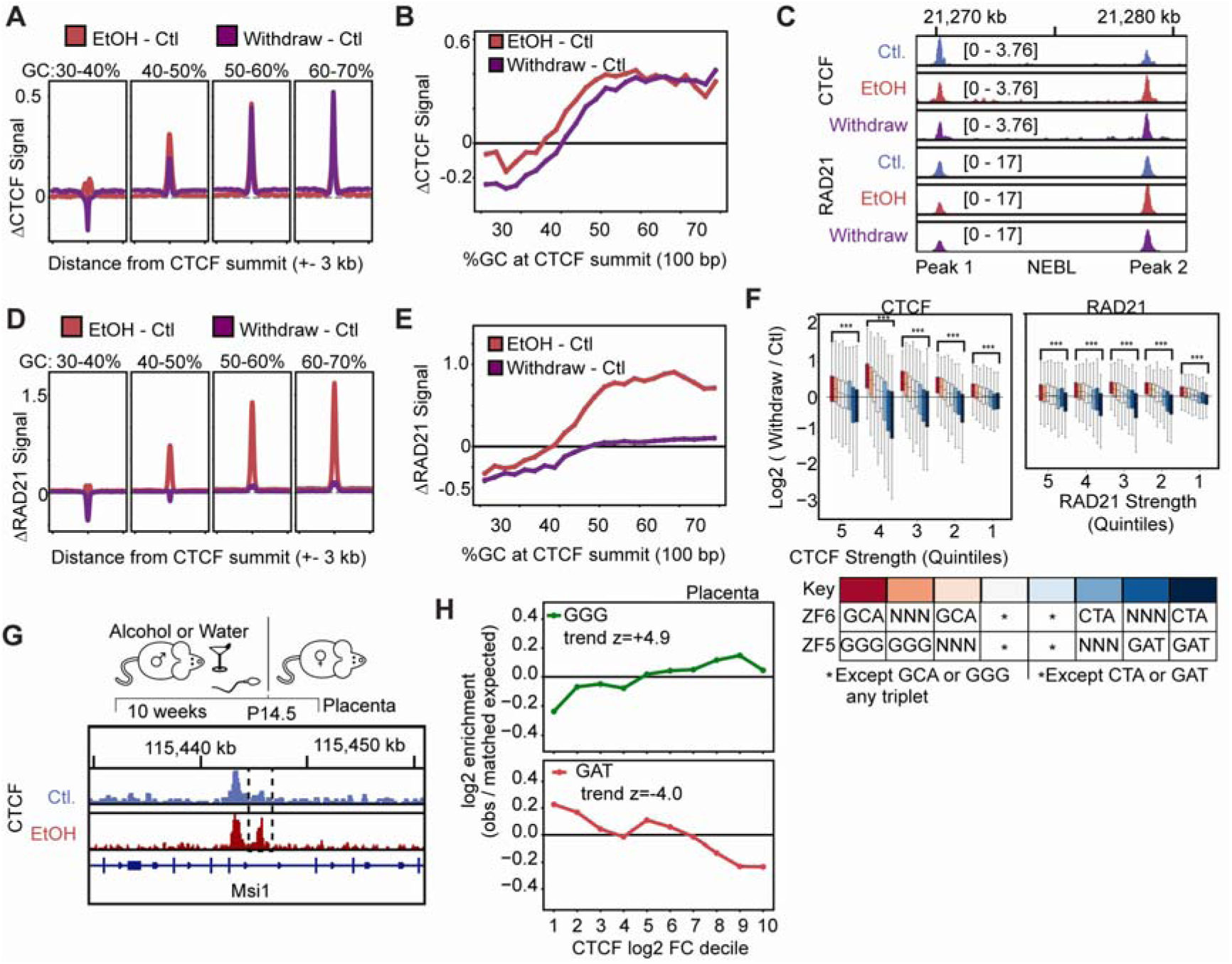
A&D) CTCF (A) and RAD21 (D) differential signal (ΔBPM) as a mean profile plot between EtOH and Withdraw conditions vs. Control at CTCF sites grouped by GC content. B&E) EtOH and Withdraw CTCF (B) and RAD21 (E) differential signal (ΔBPM) ordered by GC content. C) CTCF and RAD21 BPM signal at a locus on the NEBL gene showcasing two sites with opposing responses to ethanol; Peak 1=CTCF and RAD21 loss without recovery, Peak 2=CTCF gain without recovery and RAD21 gain with recovery. F) CTCF and RAD21 binding strength difference in withdraw condition at CTCF sites with zinc finger 6 and 5 as ‘GCA’, ‘CTA’ or ‘GGG’, ‘GAT’ respectively across binding strength quintiles. *Any triplet excluding the triplets being measured. G) Example locus in mouse placenta in both untreated and offspring with paternal ethanol exposure demonstrating an increased CTCF peak (dashed line) after paternal ethanol exposure^40^. H) Enrichment of ZF5 ‘GGG’ and ‘GAT’ by CTCF log2 fold change decile in placentas of offspring after paternal alcohol exposure. ***Wilcoxon-rank test p < 0.001

Given the persistence of CTCF changes, we expected RAD21 occupancy changes to match. Instead, RAD21 occupancy largely recovered after withdrawal, even at high-GC sites where CTCF occupancy persists (Fig. 4D-E). Investigation of the sequence motif found that while RAD21 mostly recovers, guanine content at ZF6-5 sequences had the least amount of recovery after withdrawal. The sequence ‘GCAGGG’ was identified at the loci with the highest remaining RAD21 changes after withdrawal, and ‘CTAGAT’ was at sites with persistent RAD21 loss after withdrawal (Fig. 4F). Despite these instances of residual RAD21, the level of persistence was minor compared to that of CTCF (compare Fig. 4B to 4E). These findings uncouple persistently altered CTCF occupancy from that of cohesin, or which RAD21 is a core structural subunit, indicating that the lasting effect of alcohol exposure is specific to CTCF rather than a general property of the chromatin architectural machinery.

After characterizing a phenomenon in our cell-line model where guanine content corresponds to persistent, alcohol-induced CTCF changes in our cell-line model, we asked whether the same rule operates in vivo. A previous study profiled CTCF in the placentae of mice after the paternal mouse consumed alcohol and found an overall increase in CTCF occupancy compared to placentas with unexposed sires ^40^ (Fig. 4G). Consistent with our cell-line results, we found that the degree of change corresponds to GGG vs GAT at the ZF5 position (Fig. 4H). Thus, the ZF5 guanine sequence rule governing alcohol’s effect on CTCF operates not only across metabolic states (Fig. 3H) but across a transgenerational, in vivo model of alcohol exposure (Fig. 4H).

### Alcohol exposure rewires enhancer-promoter interactions, but not strong, punctate CTCF loops

Given the substantial changes to CTCF, RAD21 and RNAPII chromatin occupancy after ethanol exposure, we tested for reciprocal changes to 3D chromatin organization. We performed in situ Hi-C in control, ethanol, and withdrawal conditions and found that, at 250 kb resolution, global patterns of 3D chromatin organization were preserved across all three conditions (Fig. 5A). Thus, alcohol does not grossly reorganize the genome at this scale.

**Figure 5:**
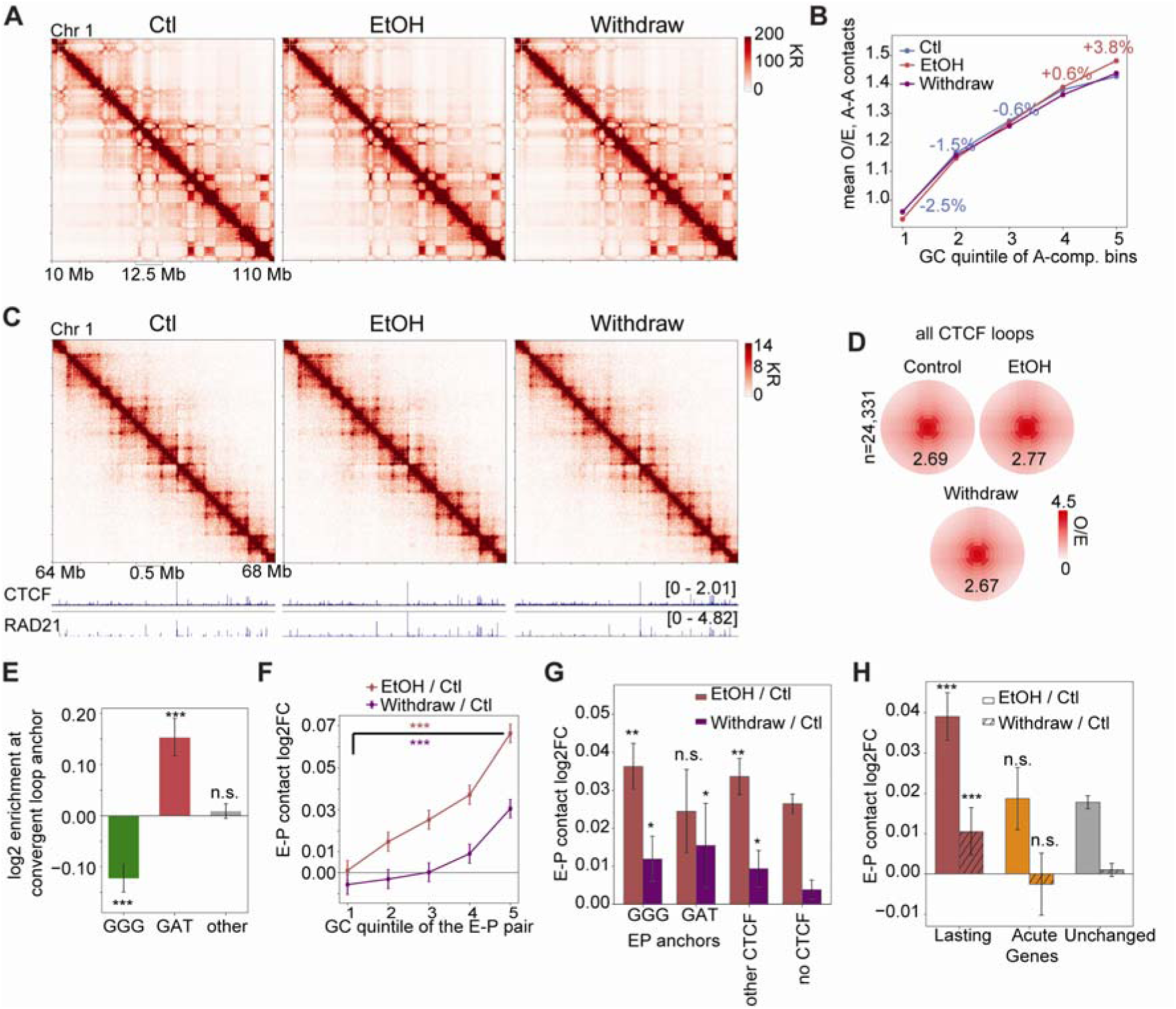
A) Hi-C contact matrix for a 100 megabase (Mb) region on chromosome 1 for control, ethanol and withdrawal conditions. KR=Knight-Ruiz matrix balanced B) Mean distance-normalized contacts between A compartments at each quintile split by GC content. C) 4 Mb contact matrix depicting chromatin loops with CTCF and RAD21 ChIP-seq tracks underneath. D) Average Hi-C signal at all CTCF loops with accompanying CTCF and RAD21 ChIP-seq profile plot at loop anchors. E) Enrichment of zinc finger 5 triplets at CTCF motifs of convergent loop anchors (n=10,039 convergent anchors). *** p<.001 Monte-Carlo permutation test, n.s.= not significant. F) Enhancer-promoter contacts (log2 (Treatment / Ctl)) by GC quintile. Error bars represent 95% confidence intervals. *** p<.001 Welch t-test of Q5 v.s. Q1. G) Enhancer-promoter contacts (log2 (Treatment / Ctl)) across CTCF sites (zinc finger 5 ‘GGG’, ‘GAT’, ‘Other’) or no CTCF. Error bars represent 95% confidence intervals. ** p <.01, * p<.05 Welch test v.s. the same condition at no CTCF sites; n.s. = not significant. H) Enhancer-promoter contacts (log2 (Treatment / Ctl)) across lasting, acute and unchanged differentially expressed genes. Error bars represent 95% confidence intervals. *** p<.001 Welch t-test v.s. each condition at unchanged genes; n.s. = not significant.

By measuring the interactions between 3D chromatin compartments, we found that compartmental interactions did not significantly increase in either active (A compartments) or inactive (B compartments) regions of the genome (Fig. S5A). Considering the impact of guanine on ethanol-mediated chromatin disruption (Fig. 3), we investigated if compartmental interactions were changed at high guanine regions and found that active compartments had slightly higher interaction intensity at guanine-rich regions, although this feature recovered in the withdrawal condition (Fig. 5B).

Surprisingly, despite the large-scale changes to CTCF and RAD21 occupancy after alcohol disruption (Fig. 3 and 4), we did not find changes to the strong, punctate CTCF loops (Fig. 5C-D). Furthermore, the contact probability curve was largely unchanged with genomic distance, including the characteristic slope at 10^5^-10^6^ bp associated with cohesin-mediated extrusion (Fig. S5B) ^41^. Even CTCF loops with a ‘GGG’ motif at the zinc finger 5 region at convergent anchors behaved akin to the rest of the CTCF loops (Fig. S5C). Investigating alcohol-sensitive CTCF binding sites further, we found that CTCF sites at convergent CTCF loop anchors are underrepresented for ‘GGG’ motifs at zinc finger 5, in contrast to ‘GAT’ which are most prevalent at loop anchors (Fig. 5E).

Because CTCF facilitates long-range DNA interactions outside of its role in forming the canonical strong CTCF loops, and since the guanine-rich CTCF binding sites most sensitive to alcohol are underrepresented at loop anchors, we next investigated the effects of alcohol on more nuanced enhancer-promoter interactions ^21^. We observed that alcohol increases global enhancer-promoter interactions as a function of guanine content, which do not fully recover after 6 days of withdrawal (Fig. 5F). CTCF-anchored enhancer-promoter interactions are also increased after alcohol exposure, with a larger increase at sites containing a ‘GGG’ motif at zinc finger 5 compared to ‘GAT’ and compared to enhancer-promoter interactions with no CTCF site at all (Fig. 5G). To connect these changes in enhancer-promoter interactions to the changes in gene expression (Fig. 1), we assessed differences in E-P interactions after alcohol and withdrawal treatments for the three groups of genes we identified (acute, delayed, lasting). While all three groups of genes have increased E-P interactions after ethanol treatment, only the ‘lasting’ subset of genes were associated with persistent changes to enhancer-promoter interactions (Fig. 5H).

These results indicate that alcohol exposure has a persistent impact on gene expression and RNA polymerase II elongation. This is accompanied by re-distribution of CTCF binding in a guanine-dependent manner and altering enhancer promoter contacts at the chromatin level, providing evidence that binge-level alcohol exposure has a lasting effect on gene expression via epigenetic disruption.

## DISCUSSION

Alcohol is metabolized to acetaldehyde, a reactive electrophile with marked specificity for guanine^25,26^ . Here, we demonstrated that this specificity converges on the guanine-rich core motif of CTCF. Alcohol exposure redistributes CTCF and cohesin across the genome along an axis defined by the guanine content of the underlying binding site, and the direction of that redistribution corresponds to the identity of individual zinc-finger triplets. This response tracks motif sequence rather than expression level or binding strength and it persists six days after alcohol is withdrawn. We therefore propose that the vulnerability of genome architecture to alcohol is encoded in the DNA sequence of the architectural binding sites themselves.

Several aspects of the chromatin response to alcohol indicate chemistry acting directly on the motif rather than a regulated signalling output. Ethanol exposure increased RNAPIIs2p at promoters while depleting it across gene bodies at every expression level (Fig. 2), a redistribution more consistent with impaired elongation through transcription-blocking lesions than with transcriptional repression^15,42^. Critically, regions that gain RNAPIIs2p were sharp, intergenic, and prominently enriched for the CTCF motif. This pattern distinguishes alcohol from the stochastic damage models established for UV, formaldehyde and oxidative stress, in which lesion burden scales with the length of the transcribed regions; here, the burden instead follows guanine content at protein-bound loci. Motif analysis defined this dependence at base-pair resolution, as GGG at the CTCF ZF5 position is enriched at sites with the largest increases in CTCF occupancy (Fig. 3). The largely additive effects across ZF positions would be difficult to reconcile with a cooperative or actively regulated mechanism and instead points to independent per-ZF chemistry at the motif. Importantly, these motif-specific effects cannot be explained by sequence alteration or cytotoxicity, as our results demonstrate a lack of mutations or change to cell-cycle/growth after alcohol exposure; thus, the effects are more consistent with an epigenetic-chemical relationship. To determine whether our findings regarding guanine-based CTCF re-distribution reflect aldehyde chemistry in general rather than a response to alcohol specifically, we assessed CTCF response to other exposure models in two independent datasets. In mouse liver, CTCF occupancy differs between fed and starved animals, conditions that differ in the production of endogenous metabolic aldehydes. Sites that gain CTCF in the fed state (aldehyde-rich) are enriched for ZF5 ’GGG’ (Fig. 3H), suggesting that the “normal” landscape of CTCF, and its relationship to guanine content (Fig. 3B, blue), may already be influenced by endogenous aldehydes. In the placentae of offspring sired by alcohol-exposed males, the increase in CTCF occupancy likewise corresponds to ’GGG’ versus ’GAT’ content at the binding site of ZF5 (Fig. 4). Thus, sequence variability at the binding site of CTCF ZF5 may serve as a predictor of response to metabolic stressors operating across metabolic states, across species and in vivo; therefore, we propose it as a general sequence-encoded aldehyde vulnerability of CTCF.

CTCF most classically shapes the 3D genome in collaboration with RAD21 to form punctate loops represented by intense signal in maps of chromatin conformation^18^. However, not all CTCF sites participate in these intense cohesin extrusion-mediated architectural loops^43^. Our withdrawal condition distinguishes this functional relationship as CTCF shows widespread persistent changes, while cohesin (RAD21) occupancy mostly returns to normal (Fig. 4). Our results further reveal these distinct functions, as altered CTCF sites are mostly absent from the anchors of the intense punctate loops but are enriched at persistently altered enhancer-promoter contacts (Fig 5). Importantly, persistent changes to CTCF-associated enhancer-promoter contacts are associated with persistent changes to gene expression, providing a potential explanation for the lasting impacts of alcohol-induced gene expression changes seen in alcohol use disorder and fetal alcohol spectrum disorder^44,45^. In total, these observations support a mechanism of chromatin memory. The CTCF binding landscape established during exposure is maintained after the removal of alcohol and is not attributable to mutation at the affected motifs.

Although our experimental design revealed important aspects of the chromatin response to cellular exposure to alcohol, some features of the design bound the scope of these conclusions and define where the model should be evaluated next. We characterized the response in a well-controlled cell line at a commonly used dose and duration; the six-day withdrawal is short relative to the timescales of human drinking and abstinence, so the persistence we observe sets a lower bound rather than an endpoint. Similarly, while the redistribution of RNAPIIs2p towards CTCF motifs led us to test occupancy changes and their relationship to guanine content, the persistence cannot be explained by DNA adducts alone as these are likely repaired before cells divide; thus, what causes CTCF changes to persist is unknown.

Taken together, our results provide a sequence-based framework for predicting which regulatory elements are vulnerable to alcohol-derived and endogenous metabolic aldehydes. Applied to the tissues and exposures relevant to alcohol-associated disease, this framework offers a mechanism linking the chemical property of aldehyde to the specific architectural and transcriptional changes that follow alcohol exposure.

## METHODS

### Alcohol Exposure Model

Cell Line studies: HCT116 cells were cultured in McCoy’s 5A Medium supplemented with 10% fetal bovine serum (FBS) and 1% penicillin / streptomycin. Ethanol was added to culture medium at a concentration of 200 mg/dL and mixed prior to adding to the cell flask. Medium was changed daily, and cells were split every third day to maintain cells in an active growing phase.

### Gene Expression (RNA-seq)

Total RNA was isolated from HCT116 cells using TRIzol reagent (Invitrogen 15596026). Ribosomal RNA was depleted with the NEBNext rRNA Depletion kit (New England Biolabs E6310), after which libraries were prepared with the NEBNext Ultra II RNA Library Prep Kit for Illumina (New England Biolabs E7770, E7775). Briefly, RNA was fragmented via incubation at 94°C, after which first strand and second strand cDNA synthesis was performed. Libraries were then generated via end prep, adaptor ligation, and PCR enrichment. Three independent biological replicates per sample were sequenced in single-end using the Illumina NovaSeq X Plus platform.

### RNA-seq Data Analysis

Single-end fastq reads were aligned to hg38 and processed using the ENCODE RNA-seq pipeline (https://github.com/ENCODE-DCC/rna-seq-pipeline) using STAR^46^ as the aligner and RSEM^47^ for transcript quantification. RSEM files were further aggregated and processed using DESeq2^48^ with a log2 fold change cutoff of 0.58 or -0.58 (1.5-fold change) and a p-adjusted value of <0.05. Processed RSEM files are available on GEO (See ‘Data Availability’). Custom python scripts used for all RNA-seq data analysis in this manuscript are available on GitHub (See Code Availability).

### Chromatin Conformation Capture (in situ Hi-C)

Two million cells per replicate were pelleted and fixed in 1% formaldehyde for 10 minutes at room temperature, quenched with 0.2 M glycine, washed, and re-suspended in Hi-C lysis buffer (Tris-HCl, 1 M, pH 8; NaCl, 2 M; Igepal CA630, 0.2%; 1X Roche protease inhibitor cocktail). After lysis, pelleted nuclei were washed and re-suspended in 0.5% sodium dodecyl sulfate for incubation at 62°C for 10 minutes. After quenching with 10% Triton X-100 and incubation at 37°C for 15 minutes, DpnII (NEB R0543M, 400 U / reaction) was added for 2 hours to digest chromatin. Next, digested chromatin was incubated at 65°C for 20 minutes to inactivate DpnII, after which time restriction fragment overhangs were marked with biotin and repaired by incubation at 37°C for one hour in the presence of fill-in master mix (15 ul of 1 mM biotin-14-dATP, 1.5 ul of 10 mM dCTP, 1.5 ul of 10 mM dGTP, 1.5 ul of 10 mM dTTP, 22.5 ul water, and 8 ul of 5 U/ul DNA polymerase I, Large (Klenow) Fragment (NEB M0210)). 900 ul ligation master mix was then added and cells incubated with slow rotation overnight (669 ul water, 120 ul of 10X NEB T4 DNA ligase buffer (NEB B0202), 100 ul of 10% Triton X-100, 6 ul of 20 mg/ml bovine serum albumin, 5 ul of 400 U/ul T4 DNA Ligase (NEB M0202)). The next day, protein was degraded by proteinase K (NEB P8107) and SDS with 55°C incubation for 30 minutes. Sodium chloride was added and chromatin incubated at 68°C overnight to reverse crosslinks. After ethanol precipitation, DNA was sheared via sonication (Diagenode Bioruptor, 18 cycles of 30 seconds on / 60 seconds off). Dynabeads MyOne Streptavidin C1 beads (Life Technologies 65602) were used to pull down biotin-containing ligated DNA fragments, and libraries prepared on the beads via end repair, A-tailing, and Illumina adaptor ligation using the NEBNext modules. Samples were sequenced in triplicate for control and ethanol samples, and 2 replicates for ethanol + withdrawal sample, with paired-end reads using the AVITI24 platform.

### Hi-C Data Analysis

Paired-end fastq files were aligned to hg38 using the juicer pipeline^49^ version 1.6. Reads were filtered by quality threshold q30 with samtools^50^ -q 30. Replicates were combined using the mega.sh script in the juicer pipeline (version 1.6). Hi-C contact matrix (.hic) files for control and ethanol treatment were subsampled using Bernoulli subsampling to 592 million contacts to match the ethanol withdrawal condition. Hi-C matrices were plotted using hic-straw, numpy and matplotlib. Chromatin loops were called using SIP^51^ at 5 kb, 10 kb and 25 kb with the following parameters: 5 kb: -mat 1000 -d 5 -res 5000, 10 kb: -mat 5000 -d 5 -res 10000, 25 kb: -mat 500 -d 2 -res 25000. All loops were rebinned to 25 kb and merged. Each annotated CTCF site at convergent loop anchors was tested for enrichment in each ZF5 class against a permutation null stratified on 12×12 grid of motif log-odds score x CTCF occupancy, in 20,000 hypergeometric draws, so that the comparison is between sites of equivalent motif strength and equivalent baseline binding.

Compartments were called with CRUSH at 10 kb on the depth-matched .hic files, using hg38 chromosome sizes, gene annotation for A seeding, and the hg38 fasta for B seeding^52,53^. For every A-A bin pair separated by 50-500 kb, the distance normalized contact value was assigned to a GC quintile defined on the mean GC of the two bins. Saddle plots were created by ranking bins into 20 quantiles genome-wide by CRUSH compartment score. Throughout the Hi-C analyses, observed contacts were divided by the genome-wide expected value at the same distance separation.

Enhancer-promoter contacts were binned at 10 kb, with promoter bins defined as those from 20 kb –2 Mb containing the TSS of a gene with TPM >1 and an enhancer defined by annotated H3K27ac. E-P pairs were quantified by the distance normalized Hi-C signal, with changes expressed as the log2 ratio. To remove any potential residual intensity dependence, contacts were MA normalized. Contacts grouped GC quintiles used the mean GC of the two bins for each pair.

Processed .hic files are available on GEO (See ‘Data Availability’). Custom python scripts used for all Hi-C data analysis in this manuscript are available on GitHub (See Code Availability).

### Protein Occupancy (ChIP-seq and CUT&Tag)

ChIP-seq: Cells were fixed with 1% formaldehyde for 10 minutes at room temperature, quenched with 0.125 M glycine, washed, and re-suspended in cellular lysis buffer (PIPES, 5 mM, pH 8; potassium chloride, 85 mM; Nonidet P40, 0.5%; 1X Roche protease inhibitor cocktail). Nuclei were collected via centrifugation and re-suspended in Sarkosinate nuclei lysis buffer (Tris-HCl, 10 mM, pH 8; NaCl, 100 mM; Sodium Deoxycholate, 0.1%; Sodium lauroyl sarkosinate, 0.5%; EDTA, 1 mM; EGTA, 0.5 mM; Triton X-100 1% (after sonication); 1X Roche protease inhibitor cocktail). After sonication, cellular debris was pelleted and supernatants containing sheared fixed chromatin were used for immunoprecipitation. A pre-washed Protein A / Protein G mixture was used to pre-clear lysates for 2 hours, after which time antibody-coated beads were added to supernatants overnight. Antibody-coated beads with bound chromatin were then washed, crosslinks were reversed at 65°C overnight, and DNA isolated via Phenol:chloroform:isoamyl alcohol extraction. Libraries were prepared via the iNext protocol and samples submitted for sequencing. Samples were sequenced in triplicate for control, ethanol and ethanol + withdrawal samples, with paired-end reads via the Illumina NovaSeq X Plus platform.

CUT&Tag: 100,000 cells per reaction were resuspended in Nuclei Extraction Buffer for 10 minutes on ice (20 mM HEPES-KOH, pH 7.9, 10 mM KCl, 0.1% Triton X-100, 20% glycerol, 7 mM spermidine, 1X Roche EDTA-Free Protease Inhibitor). Nuclei were centrifuged and incubated with 10 uL activated ConA beads for 10 minutes at room temperature. After binding, beads were incubated with primary antibody overnight (RNApol2-Ser2, Cell Signaling E173G). Secondary antibody was added the next day for 30 minutes at room temperature, then tagmentation completed after washing via addition of pAG-Tn5 in the presence of buffer containing 10 mM MgCl2 for enzyme activation. Reactions were quenched and tagmented chromatin fragments were released into solution in the presence of SDS buffer, followed by PCR amplification. PCR amplification on isolated tagmented DNA fragments was completed with unique i5 and i7 primers for each sample. Samples were sequenced in triplicate with paired-end reads via the Illumina NovaSeq X Plus platform.

### Protein Occupancy Data Analysis

ChIP-seq samples were processed using the ENCODE ChIP-seq pipeline (https://github.com/ENCODE-DCC/chip-seq-pipeline2). All ChIP-seq samples passed the ENCODE replicate concordance standard. A minimum quality threshold of 30 was used for all RAD21 and CTCF samples. Peaks were called with macs2^54^, with downstream peak analysis done using the conservative IDR set of peaks over all replicates for that condition. Differential peaks were called using MAnorm^55^ with a window size of 1000 for both RAD21 and CTCF. Bigwigs were generated using bamCoverage^56^ -- binSize 10 -- normalizeUsing BPM -- extendReads to normalize for sequencing depth. Technical bias and variation were normalized in the bigwigs by applying the MA normalization to all genomic bins.

CTCF and RAD21 motif analysis was carried out by obtaining called peaks with macs2 selecting peaks with only a single CTCF motif within +/- 150 bp of the summit (macs2 --call-summits). This was achieved using bedtools^57^ getfasta for each 300 bp window, followed by FIMO^58^ motif finder with the JASPAR^59^ CTCF motif. Only peaks with a single motif found were kept for further analysis. The motif was then used for CTCF peak orientation and then subdivided into zinc finger motifs in the 19 bp core motif as each zinc finger corresponds to a nucleotide triplet e.g. zinc finger 5: positions 10,11,12. Sites were ranked by log2 fold change in occupancy and split into deciles (D1: most lost to D10: most gained) and enrichment of a given ZF5 triplet within each decile was assessed against a stratified permutation null: sites cross-classified into 10×10 grid by motif log-odds score decile and by mean peak intensity decile, and within each grid position the triplet-positive sites were redistributed across the fold-change deciles by a multivariate hypergeometric draw, 20,000 permutations. Per decile we report the observed count divided by the permutation mean (log2 scale) with an overall monotonic-trend statistic expressed as a z-score against its permutation distributions.

Mouse liver (fed vs 24-h starved) CTCF peaks were obtained from the IDR optimal peak sets^39^. Peak summits from the two conditions were merged into union sites by greedy clustering within 250 bp, keeping the highest signal summit per cluster. Reads were counted over +/- 250 bp of each summit, converted to CPM, and log2 fold change computed as fed over starved, with sites below a minimum coverage of 20 reads excluded. Mouse placenta (paternal ethanol vs control) CTCF occupancy used a similar pipeline, but with MAnorm performing the peak union clustering. Zinc-finger triplet assignment and enrichment testing for both datasets followed the human cell-line procedure described above. CUT&Tag fastq reads were aligned to hg38 with bowtie2 using the following parameters: --end-to-end, -- very-sensitive, --no-mixed, --no-discordant, --phred33 -I 10 -X 700 and further filtered with samtools -q 10. Bigwig files were created using bamCoverage --normalizeUsing BPM. Peak calling was performed using macs3 with the -f BAMPE parameter. Heatmaps and profile plots were created using computeMatrix and plotHeatmap from deeptools. Processed bigwig and broadPeak files are available on GEO (See ‘Data Availability’). Guanine content was measured as GC percentage using the UCSC hg38 5-bp GC track, taken as the mean over a 100 bp window centered on ChIP-seq summits. Peak summits were assigned to bins based on GC content, each sorted by GC content within the bin. Signal matrices were built with deepTools computeMatrix, and differential matrices were computed bin-wise for EtOH - Ctl or Withdrawal - Ctl from MA-normalized BM matrices. Custom python scripts used for all ChIP-seq and CUT&Tag data analysis in this manuscript are available on GitHub (See Code Availability).

### Mutation Analysis

Somatic single-nucleotide variants were called from CTCF ChIP-seq reads across control and ethanol conditions using Mutect2^60^ with the 6-day alcohol condition as the “tumor” sample and control as the matched normal, using the hg38 reference genome and gnomAD^61^ as a germline population prior. Read orientation bias was modeled with LearnReadOrientationModel and applied during statistical filtering with FilterMutectCalls, and only PASS variants were retained. Calls were restricted to biallelic SNPs to avoid the ambiguity in allele-frequency estimation introduced by multi-allelic sites, and variants overlapping the ENCODE hg38 blacklist (v2)^62^ were excluded, yielding 1,899 candidate variants. Putative variants were intersected with a master list of CTCF peaks irrespective of condition and CTCF peaks were classified as gained, lost, or unbiased in alcohol based on log2 fold change in occupancy (>1, <-1, or intermediate, respectively). A permutation test (10,000 iterations) was used to compare candidate-variant density between gained, lost, and unbiased CTCF sites, and allele frequencies of candidate variants were plotted. Performed analyses are available (See ‘Code Availability’).

### Antibodies

ChIP-seq: CTCF Abcam ab70303, RAD21 Abcam ab992. CUT&Tag: Phospho-Rpb1 CTD (Ser2) E1Z3G Cell Signaling.

### Flow Cytometry

Cell cycle analysis was assessed by fixing one million cells with 70% ethanol on ice for one hour, washing cells with PBS, and then re-suspending in staining buffer containing 100 ug/mL RNase A, 50 ug/mL Propidium Iodide, and 0.1% Triton X-100. After an overnight incubation at 4°C, cell cycle phase was obtained using a BD FACSCalibur flow cytometer.

## DATA AVAILABILITY

All genomics data generated for this study have been deposited in the GEO database under the following accession codes; RNA-seq (<u>GSE342175</u>), CUT&Tag RNA polymerase II serine 2 phosphorylation (<u>GSE342529</u>), ChIP-seq CTCF and ChIP-seq RAD21 (<u>GSE341870</u>), Hi-C (<u>GSE342173</u>). Previously published data from placental CTCF occupancy after paternal alcohol exposure^40^ used in figure 4 (GEO accession: <u>GSE203189</u>). Metabolic regulation of CTCF chromatin occupancy from starvation^39^ used in figure 3 (GEO accession: <u>GSE207692</u>). The 18 ENCODE transcription factor binding sites used in Figure 2 are available at the following ENCODE accessions: CTCF (<u>ENCFF832INR</u>), CBX3 (<u>ENCFF423XAH</u>), CEBPB (<u>ENCFF963FVR</u>), EGR1 (<u>ENCFF822VKJ</u>), ELF1 (<u>ENCFF420PYW</u>), FOSL1 (<u>ENCFF936WPZ</u>), JUND (<u>ENCFF378LWF</u>), MAX (<u>ENCFF214KLO</u>), RAD21 (<u>ENCFF391AAM</u>), SIN3A (<u>ENCFF493SWH</u>), SP1 (<u>ENCFF600MKH</u>), SRF (<u>ENCFF502RNC</u>), TCF7L2 (<u>ENCFF845TYG</u>), TEAD4 (<u>ENCFF008LFE</u>), USF1 (<u>ENCFF725PAR</u>), YY1 (<u>ENCFF451EDW</u>), ZBTB33 (<u>ENCFF141RIH</u>), ZFX (<u>ENCFF858YWR</u>).

## CODE AVAILABILITY

All custom analyses produced in this manuscript were completed using a combination of R, Python and the Linux command line. Scripts used to generate the graphics and analyses have been stored on the following GitHub repository, enabling open access for reproduction of analysis ( https://github.com/BenNolann/AlcoholChromatin )

## ACKNOWLEDGEMENTS

We thank Dr. Padmashri Ragunathan, Dr. Adam Karpf, Dr. Shibiao Wan, and Dr. Ryan Wong for continued support and advice throughout the duration of this project. We thank the Flow Cytometry Research Core Facility at the University of Nebraska Medical Center for assistance with cell cycle analysis. We are grateful to the authors of the previously published datasets reanalyzed here, and to the ENCODE Consortium and the ENCODE production laboratories for generating the transcription factor binding data used in this study.

## AUTHOR CONTRIBUTIONS

B.N. - Conceptualization, formal analysis, writing: original draft, editing, Illustration; J.A.P, I.M.QP., S.M., R.A.W., A.S., L.J., C.O. – Methodology; T.E.R, A.E., H.L.H., - Writing: review and editing; C.T.C. – Conceptualization, methodology, writing: review and editing, funding acquisition, resources; M.J.R. Conceptualization, formal analysis, writing: review and editing, funding acquisition, resources.

