## Supplemental Figures for "Guanine content in the CTCF motif encodes a vulnerability to persistent, alcohol-induced redistribution"

**SUPPLEMENTARY DATA**

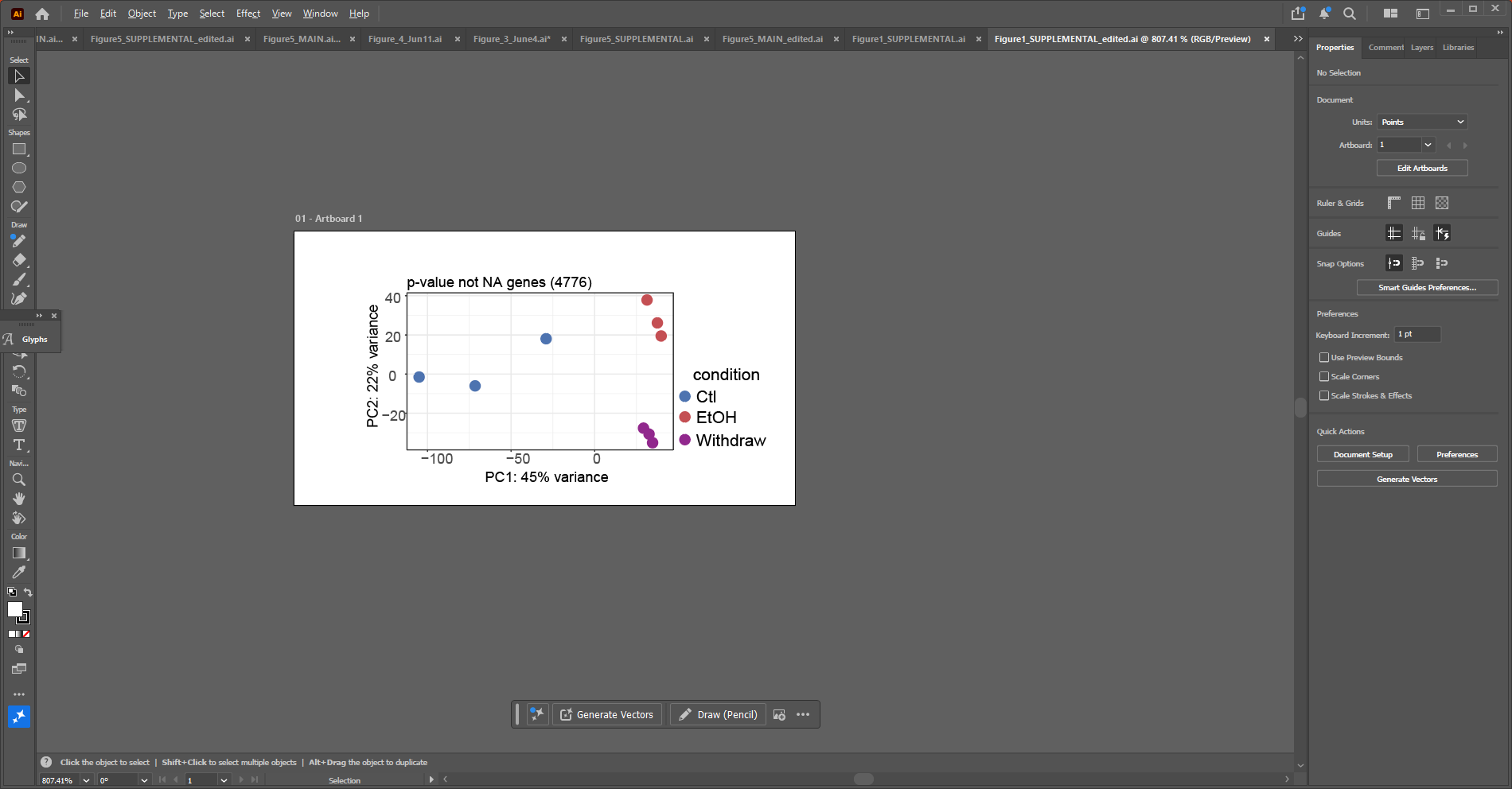

**Supplementary Figure S1:** a) PCA for genes with a non-NA adjusted p-value. These genes are used for downstream differential gene expression analysis.

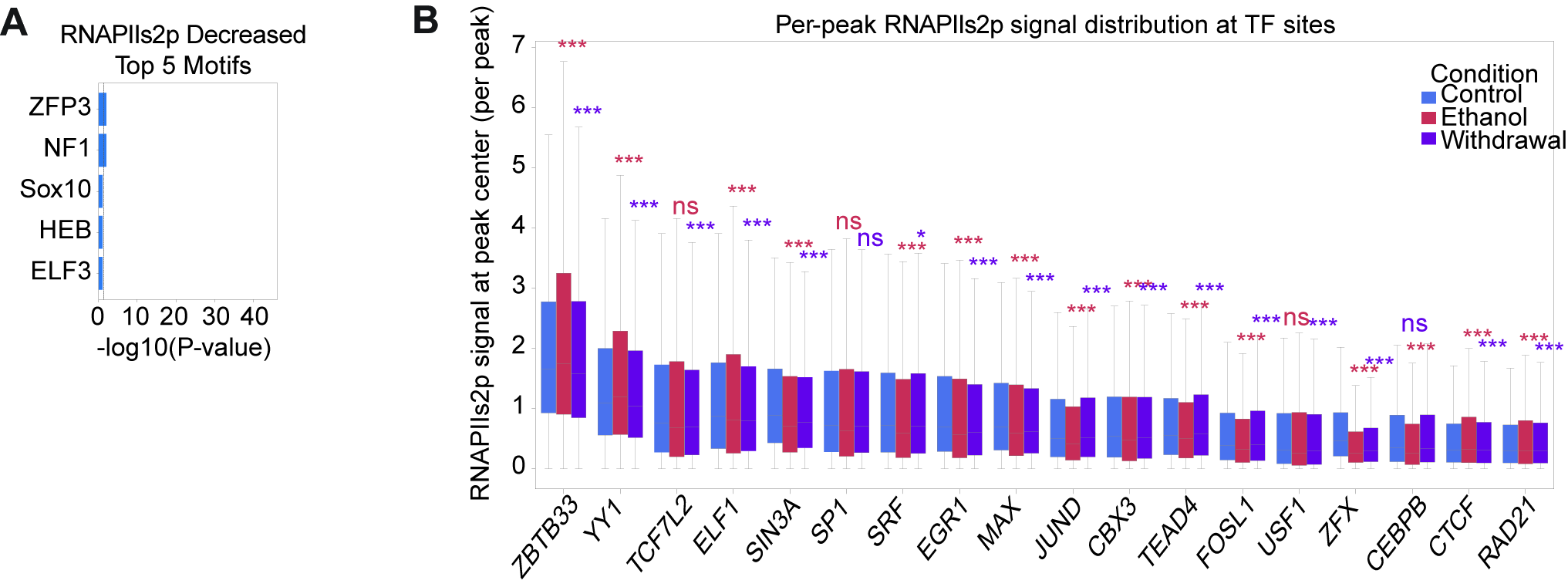

**Supplementary Figure S2**: A) Top 5 motifs via Homer known results B) Mean RNAPIIs2p signal distribution from peak center +/- 50 bp. Wilcoxon rank test was performed for either Ethanol or Withdrawal versus Control for each of the 18 TFs. All statistics can be seen in Table S2.

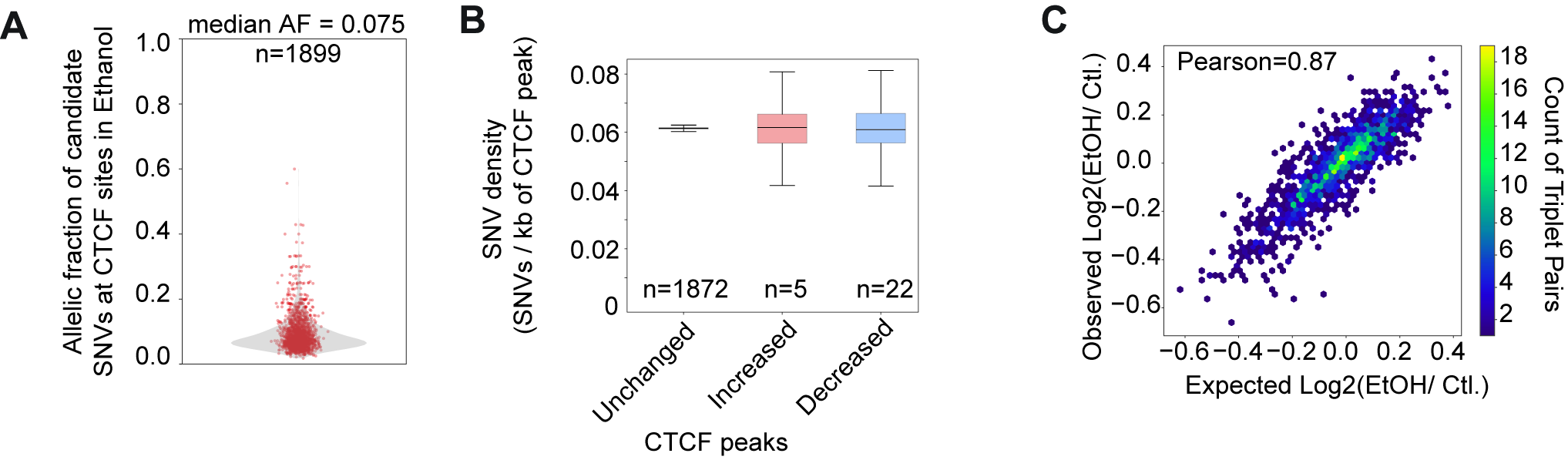

**Supplementary Figure S3:** A) Allelic fraction of SNVs called in sequencing data after ethanol exposure. B) Number of SNVs per kb at CTCF peaks that are unchanged, increased, or decreased after ethanol exposure. C) Density plot of CTCF binding sites disruption after alcohol grouped by CTCF motifs, paired by zinc finger triplets (i.e. ZF5=GGG and ZF6=GCA). Observed CTCF disruption is the actual log2 ( EtOH / Ctl ) of CTCF sites with a specific pair of Zinc Finger motifs. Expected CTCF disruption is the addition of CTCF sites with each of two zinc finger motifs separately.

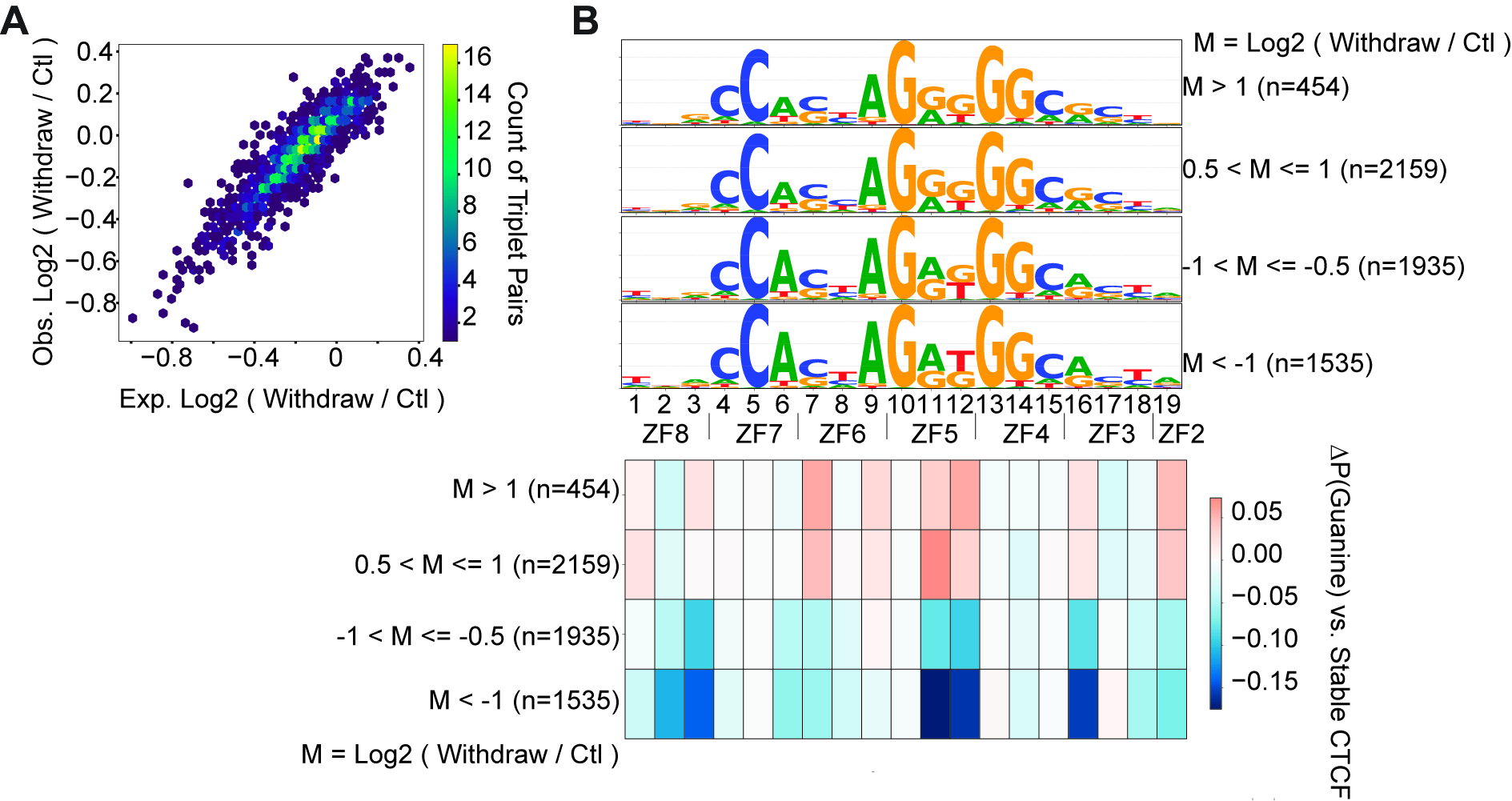

**Supplementary Figure S4:** A) Density plot of CTCF binding sites disruption after withdrawal from alcohol grouped by CTCF motifs, paired by zinc finger triplets (i.e. ZF5=GGG and ZF6=GCA). Observed CTCF disruption is the actual log2 ( Withdraw / Ctl ) of CTCF sites with a specific pair of Zinc Finger motifs. Expected CTCF disruption is the addition of CTCF sites with each of two zinc finger motifs separately. B) top: CTCF average logo for each group of binding disruption after withdrawal condition in four groups of varying susceptibility. Bottom: Heatmap of the difference in guanine probability at each nucleotide of the CTCF binding in four groups of varying alcohol withdrawal susceptibility.

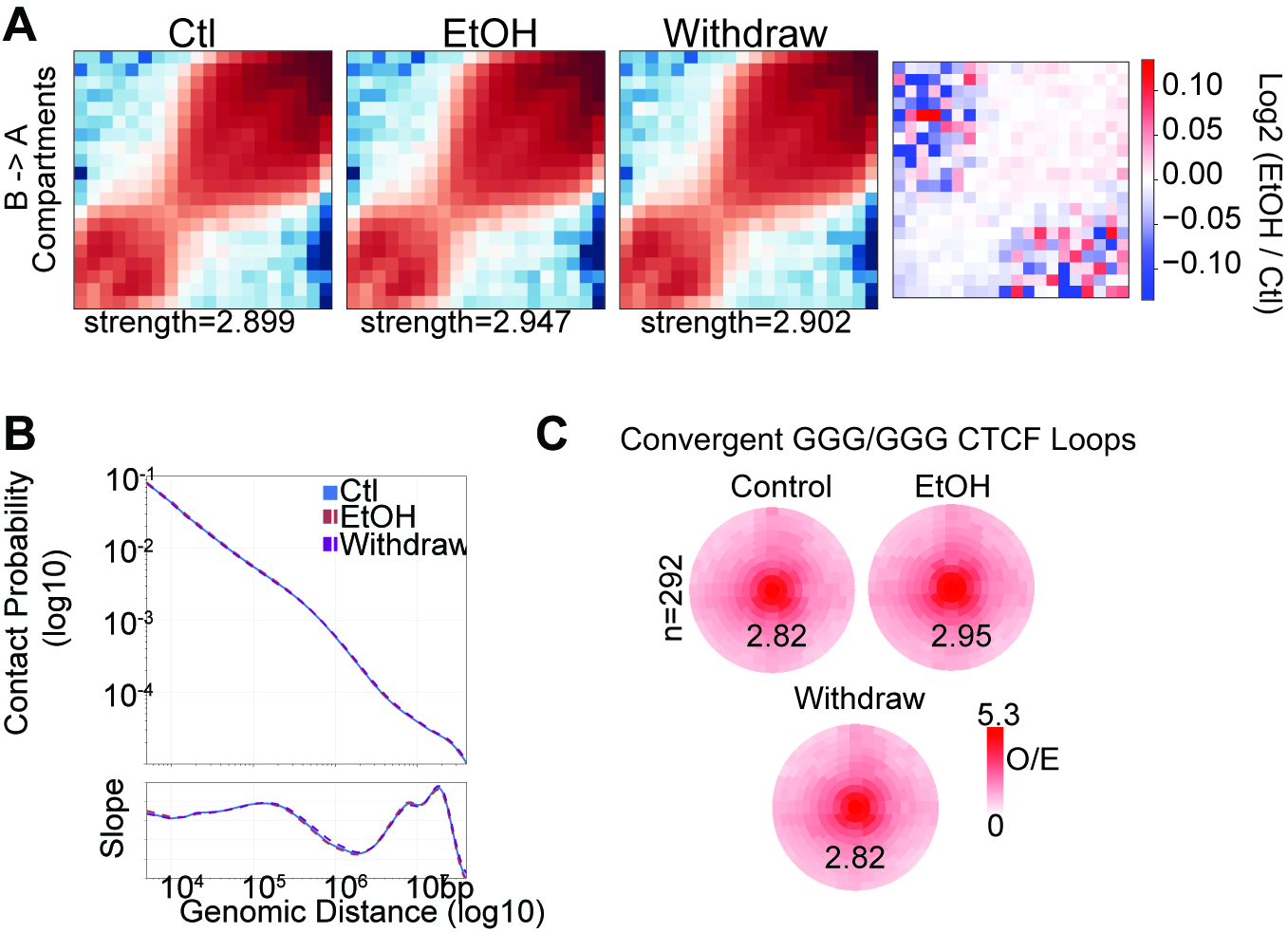

**Supplementary Figure S5:** A) Saddle plot for compartment values from Ctl, EtOH, Withdraw conditions with a difference plot of EtOH against Ctl. B) Contact probability of all three Hi-C matrices. C) CTCF loops with convergent oriented CTCF sites that contain GGG at the ZF5 position at both anchors, and the signal in Control, Ethanol, and Withdrawal conditions.

|  | control |  |  | ethanol |  |  |
| --- | --- | --- | --- | --- | --- | --- |
|  | rep1 | rep2 | rep3 | rep1 | rep2 | rep3 |
| Day 1 | 10.65 | 7.55 | 5.66 | 6.35 | 7.3 | 5.2 |
| Day 2 | 21.25 | 10.95 | 30.5 | 21.05 | 25.8 | 27.1 |
| Day 3 | 34.45 | 29.05 | 35.6 | 40.3 | 27.9 | 37.65 |
| Day 4 | 8.85 | 8.4 | 5.8 | 12.55 | 7.55 | 4.65 |
| Day 5 | 23.6 | 17.1 | 8.4 | 22.45 | 18.6 | 10.5 |
| Day 6 | 26.25 | 22.4 | 26.85 | 40.4 | 22.5 | 20.35 |

**Supplementary Table S1:** Viable cells per replicate in control or each day during ethanol treatment.

| **TF** | **Comparison** | **n_peaks** | **median_diff** | **wilcoxon_p** | **padj** | **sig** |
| --- | --- | --- | --- | --- | --- | --- |
| CEBPB | Ethanol vs Control | 17686 | -0.052349 | 0 | 0 | *** |
| CTCF | Ethanol vs Control | 52578 | 0.019599409 | 0 | 0 | *** |
| ZFX | Ethanol vs Control | 23434 | -0.133008545 | 0 | 0 | *** |
| YY1 | Ethanol vs Control | 18546 | 0.1022185 | 9.53E-289 | 4.29E-288 | *** |
| FOSL1 | Ethanol vs Control | 26698 | -0.041419955 | 1.99E-243 | 7.17E-243 | *** |
| JUND | Ethanol vs Control | 24426 | -0.052506818 | 4.18E-240 | 1.25E-239 | *** |
| SIN3A | Ethanol vs Control | 19975 | -0.102406364 | 5.20E-238 | 1.34E-237 | *** |
| RAD21 | Ethanol vs Control | 51153 | 0.002156364 | 3.87E-184 | 8.71E-184 | *** |
| ZBTB33 | Ethanol vs Control | 5817 | 0.129136636 | 2.84E-75 | 5.68E-75 | *** |
| SRF | Ethanol vs Control | 10412 | -0.058345955 | 1.09E-54 | 1.96E-54 | *** |
| MAX | Ethanol vs Control | 33689 | -0.041252364 | 4.57E-54 | 7.47E-54 | *** |
| TEAD4 | Ethanol vs Control | 6876 | -0.035613682 | 5.74E-30 | 8.60E-30 | *** |
| EGR1 | Ethanol vs Control | 15041 | -0.045639727 | 4.76E-24 | 6.59E-24 | *** |
| CBX3 | Ethanol vs Control | 28669 | -0.012267273 | 2.27E-10 | 2.92E-10 | *** |
| ELF1 | Ethanol vs Control | 5380 | 0 | 2.31E-06 | 2.77E-06 | *** |
| SP1 | Ethanol vs Control | 30628 | -0.016516727 | 0.159418671 | 0.179346005 | ns |
| TCF7L2 | Ethanol vs Control | 19159 | -0.010080455 | 0.284884064 | 0.30164195 | ns |
| USF1 | Ethanol vs Control | 10617 | 0 | 0.616320203 | 0.616320203 | ns |
| ZFX | Withdrawal vs Control | 23434 | -0.126352909 | 0 | 0 | *** |
| SIN3A | Withdrawal vs Control | 19975 | -0.078834 | 5.43E-231 | 4.89E-230 | *** |
| MAX | Withdrawal vs Control | 33689 | -0.045166364 | 7.82E-179 | 4.69E-178 | *** |
| EGR1 | Withdrawal vs Control | 15041 | -0.060345818 | 5.36E-151 | 2.41E-150 | *** |
| TCF7L2 | Withdrawal vs Control | 19159 | -0.033978727 | 7.62E-84 | 2.74E-83 | *** |
| RAD21 | Withdrawal vs Control | 51153 | 0 | 4.70E-37 | 1.41E-36 | *** |
| YY1 | Withdrawal vs Control | 18546 | -0.022662136 | 1.97E-15 | 5.06E-15 | *** |
| USF1 | Withdrawal vs Control | 10617 | -0.004164909 | 7.02E-14 | 1.58E-13 | *** |
| ELF1 | Withdrawal vs Control | 5380 | -0.024543955 | 2.03E-13 | 4.06E-13 | *** |
| FOSL1 | Withdrawal vs Control | 26698 | 0 | 3.32E-11 | 5.43E-11 | *** |
| ZBTB33 | Withdrawal vs Control | 5817 | -0.044530455 | 3.14E-11 | 5.43E-11 | *** |
| CTCF | Withdrawal vs Control | 52578 | 0 | 8.83E-10 | 1.32E-09 | *** |
| TEAD4 | Withdrawal vs Control | 6876 | 0.006233045 | 2.60E-08 | 3.60E-08 | *** |
| JUND | Withdrawal vs Control | 24426 | 0 | 1.20E-05 | 1.55E-05 | *** |
| CBX3 | Withdrawal vs Control | 28669 | -0.000450545 | 0.000615275 | 0.00073833 | *** |
| SRF | Withdrawal vs Control | 10412 | -0.005164045 | 0.037888157 | 0.042624176 | * |
| CEBPB | Withdrawal vs Control | 17686 | 0 | 0.064659447 | 0.068462944 | ns |
| SP1 | Withdrawal vs Control | 30628 | -0.000170409 | 0.414947805 | 0.414947805 | ns |

**Supplementary Table S2:** RNAPIIs2p BPM signal at center +/- 50 bp for 18 transcription factor binding sites. These data are visualized in Figure S2d.
